# ArtiDock: accurate Machine Learning approach to protein-ligand docking optimized for high-throughput virtual screening

**DOI:** 10.1101/2024.03.14.585019

**Authors:** Taras Voitsitskyi, Ihor Koleiev, Roman Stratiichuk, Oleksandr Kot, Roman Kyrylenko, Illia Savchenko, Vladyslav Husak, Semen Yesylevskyy, Sergii Starosyla, Alan Nafiiev

## Abstract

Classical protein-ligand docking has been a cornerstone technique in computational drug discovery for decades, but has reached an accuracy and performance plateau. Recently introduced Machine Learning (ML) based docking methods offer a promising paradigm shift, but their practical adoption is hampered by accuracy-to-speed trade-offs, inadequate benchmarking standards, and questionable chemical validity of predicted poses. In this study, we introduce ArtiDock - an ML-based docking technique optimized for high-throughput virtual screening applications. To evaluate ArtiDock, we developed a dedicated performance and accuracy benchmark for pocket-specific rigid protein-ligand docking, which mimics realistic industrial drug discovery scenarios and is based on the novel PLINDER dataset. We demonstrate that ArtiDock is 29-38% more accurate in comparison to leading open-source and commercial classical docking techniques such as AutoDock, Vina, and Glide, while providing a low computational cost. ArtiDock notably excels in challenging docking scenarios involving unbound protein structures and binding sites containing ions and structured water molecules. Our results show that ArtiDock could be considered as a method of choice in high-throughput virtual screening scenarios.

## Introduction

Classical or physics-based docking has been one of the foundational tools in computational drug discovery for decades. During this extended period, the docking technology has witnessed an impressive scaling up and a universal adoption in academia and industry, while remaining remarkably unchanged in terms of algorithms used. Indeed, all the major docking scoring functions have not evolved much since their introduction. All inherent limitations of classical docking, such as poor handling of the metal atoms, coordination bonds, polarization, charge transfer, and entropic contribution from water, persist to date with no clear trend for improvement. There is currently an implicit consensus in the community that classical docking has reached the practical plateau of accuracy, and no significant progress could be made without a paradigm change [1], [2].

Such a change has emerged in recent years with the appearance of the ML techniques for ligand pose prediction, often colloquially called an “AI docking”. In contrast to the classical docking, which is based on minimization of some physics-based scoring function, these techniques leverage a completely data-centric approach by learning from experimentally determined protein-ligand complexes.

The first generation of the ML docking techniques, such as DeepDock [3], TANKBind [4], EquiBind [5], and Uni-Mol [6], demonstrated results that were subpar to conventional docking in terms of accuracy and chemical validity [7], while being significantly faster. More heavyweight tools, such as the diffusion model DiffDock [8], [9], or cofolding methods RoseTTAFold All-Atom [10], NeuralPLexer3 [11], AlphaFold3 [12], Chai-1 [13], Boltz-1 [14], and Protenix [15], have shown an impressive boost in accuracy that comes, however, at the expense of very complex architectures, large model sizes, and slow training and inference.

However, it quickly became evident that accurate and unbiased assessment of ML docking techniques remains challenging due to the absence of commonly accepted, reliable benchmarks suitable for these data-driven methods. The majority of current evaluations overlook the data leakage between training and test datasets, which can result in overly optimistic performance metrics [16], [17]. In particular, the performance of cofolding tools, often showing the greatest accuracy, appears to be strongly correlated with the similarity between their test and training molecular structures [17].

As a result, existing ML docking tools are unable to replace the classical ones in the practical high-throughput virtual screening tasks for two main reasons: an unfavorable accuracy-to-speed ratio and the absence of proper evaluation for ML techniques. In other words, we still have not reached the much-anticipated docking paradigm shift.

The recently released PLINDER dataset [18] addresses most of the existing benchmarking challenges. This dataset contains a large and diverse predefined training subset from the Protein Data Bank (PDB) [19]. The train-test splitting process is cluster-based, which minimizes both the data leakage between the sets and the test set redundancy. The split considers both protein sequence similarity and protein-ligand interaction similarity, while prioritizing the inclusion of biologically relevant, high-quality structures in the test [20].

In this work, we evaluated the latest version of our proprietary ML-based protein-ligand docking tool, ArtiDock (Figure 1), against the leading open-source and commercial classical docking programs such as AutoDock4 (referred to as AutoDock-CPU hereafter) [21], [22], AutoDock-GPU [23], Vina [24], [25], and Glide [26], [27], [28], [29]. Our primary goal was to evaluate these techniques in the context of high-throughput rigid docking into a known pre-defined protein binding pocket, which reflects the most common use case in real-world drug discovery projects.

**Figure 1.**
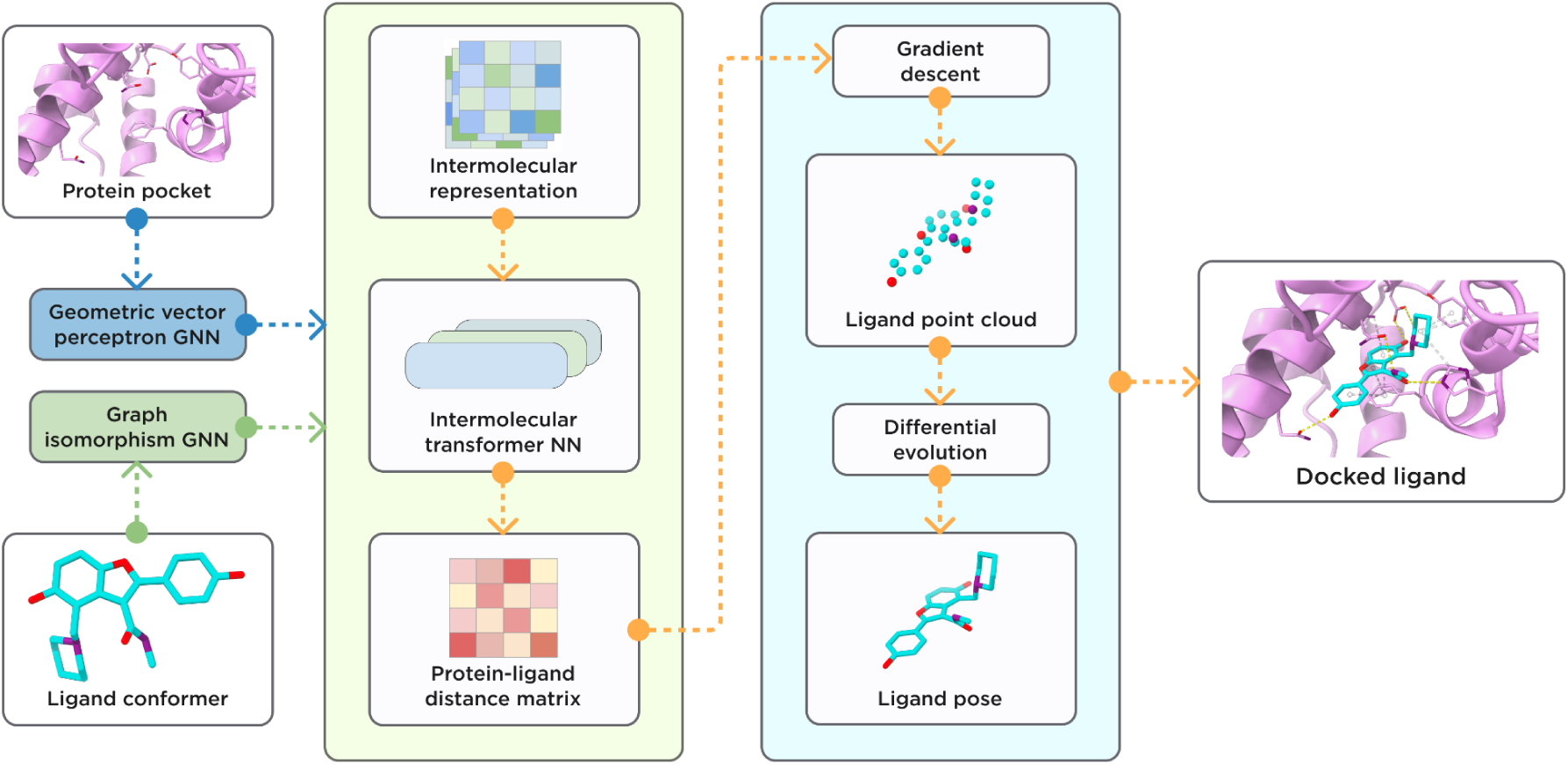
General scheme of ArtiDock model architecture and inference pipeline.

To ensure an unbiased evaluation of ArtiDock, we re-trained the model on the PLINDER training split and assessed its final performance, alongside competitors, on the PLINDER test set. The evaluation covers docking accuracy, chemical and geometric validity of the predicted poses, and computational efficiency. The test cases also represented scenarios when the input protein structure is predicted or obtained from the ligand-unbound state, as well as cases with organic cofactors, ions, or water molecules in the binding pockets.

In most comparisons between traditional and ML-based docking methods, default suboptimal parameters are used for the classical tools, or parameter selection doesn’t consider computational cost [7], [30]. This is an important omission, as two techniques with comparable accuracy may have very different throughput and thus different utility in real-world applications. To address this, we conducted a series of experiments to optimize traditional docking methods for the trade-off between accuracy and runtime, ensuring a fair and practically relevant setup for evaluation.

Our results demonstrate that ArtiDock considerably outperforms all tested physics-based methods in terms of accuracy while offering throughput on par with the fastest traditional tools. Thus, our ML docking approach pretends to be a method of choice in high-throughput virtual screening applications.

In this paper, we describe ArtiDock architecture, inference pipeline, training data preparation, and training procedure; outline the selection of optimal classical docking parameters; provide a comprehensive performance comparison and discuss the limitations and future directions.

## Methods

### Evaluation datasets

For the classical docking parametrization experiments, we used the PoseBusters Benchmark set [7] that consists of 308 crystal complexes from the PDB, each representing a unique protein-ligand pair (PLP). We removed 2 PLPs that failed to be processed by the evaluation utility.

The test split of the PLINDER dataset (release 2024-06, iteration v2) [18] was selected for the final benchmarking of classical docking approaches and ArtiDock. After the initial preprocessing and quality filtering (Appendix A), the test data contained 1,035 PLPs with 676 unique ligand Chemical Component Dictionary (CCD) codes. We retained a single ligand per CCD that forms the maximum number of non-covalent interactions with the protein. As a result, the final PLINDER subset consisted of 676 PLPs from 660 systems representing 656 PDB entries. We removed one PLP that failed to be processed by the evaluation utility. All the non-protein entities (heteroatoms) were kept in the test systems to assess the methods’ performance on the pockets containing cofactors, ions, and water molecules.

We also used a subset of PLINDER test systems proposed in the Machine Learning in Structural Biology (MLSB) 2024 challenge [31]. The PLINDER MLSB test comprises 346 single ligand-single protein systems. The main features of this dataset are unbound input protein structures from another system (apo conformations) and protein structures predicted by ML methods [20]. This dataset represents a realistic docking scenario in which the exact bound (holo) conformation of the protein is unknown. To ensure consistent pocket definitions across both unbound and bound structures, we superimposed the apo/predicted proteins onto their holo counterparts with the MolAR molecular modeling library [32], aligning the ligand-binding sites, defined as all residues within 8Å of the resolved ligand. After the preprocessing, the final PLINDER MLSB test set contained 344 PLPs.

As we re-trained the ArtiDock on the PLINDER train split, the data leakage between the splits is kept at a minimum, which avoids potential overfitting to any test samples and ensures objective model capabilities evaluation. However, this prevents us from including other ML-based docking techniques in the comparison because they also have to be re-trained on the same split, which is not technically feasible.

### Evaluation metrics

To evaluate the predicted ligand poses, we used three types of scores:

- Symmetry-Corrected Root Mean Square Deviation (RMSD). This score represents absolute deviation between experimentally determined and predicted ligand poses [33].
- Local Distance Difference Test for Protein-Ligand Interactions (lDDT-PLI). This score characterizes the extent of reproduction of the protein-ligand interactions present in the experimental complexes [33].
- PoseBusters quality checks, designed to measure the physical and chemical correctness of predicted ligand poses [7].

The scores were calculated with the PLINDER Python API that, under the hood, used OpenStructure v.2.8.0 [34], [35] and PoseBusters v.0.3.1 [7] packages.

For the benchmarking, we selected the top-ranked predicted pose based on the internal scoring function of each technique and reported the median RMSD, fraction of samples with RMSD < 2Å, median lDDT-PLI, fraction of samples with lDDT-PLI > 0.5, and fraction of samples passing all PoseBusters quality checks (PB-Valid).

### Classical docking setup

#### AutoDock-based methods

The AutoDock-CPU, AutoDock-GPU, and Vina require input protein and ligand in a PDBQT format. The same format is used to represent output ligand poses. Therefore, we applied a unified pre-/postprocessing pipeline for these docking tools.

The ligand preparation and post-docking processing were performed using the Python API of Meeko v.0.6.1 [36]. For the AutoDock-CPU and AutoDock-GPU, ligand macrocycles were set to be rigid to avoid chemically invalid structures in the output.

The protein PDBQT files were obtained with MGLTools v.1.5.7 [37] using options to protonate molecules and merge charges from non-polar hydrogens and lone pairs. We noticed that this tool fails to assign charges to water molecules and some metals. To rectify that, we added common oxidation states for these entities [38] in the final files.

Note that there were some differences in the protein and ligand preparation for docking parameterization experiments and for the final PLINDER benchmark described in Appendix B.

For the AutoDock-CPU and AutoDock-GPU, grid parameter files (GPF) were created with MGLTools, and AutoGrid v.4.2.6 [39] was used for the grid generation.

The AutoDock-CPU docking was launched with AutoDock v.4.2.6 [39] using an MGLTools-generated docking parameter file (DPF).

The AutoDock-GPU v.1.6 docking was compiled from source [40] with DEVICE=GPU, and NUMWI=256.

The Vina v.1.2.6 [41] Python API was used to compute affinity maps and perform docking.

Due to the random variability of AutoDock-CPU, AutoDock-GPU, and Vina outputs between the launches, all the results are reported as the average of three independent docking runs.

#### Glide

We used Schrödinger Suite v.2024-3 command-line interface to prepare the input data and to launch the Glide docking.

The ligand molecules were processed with the LigPrep tool and default parameters. Proteins were prepared with PrepWizard, explicitly retaining all the water molecules in the processed protein structures.

### Classical docking parametrization

To obtain an optimal docking setup in terms of time/quality tradeoff, we launched a range of experiments with varying docking parameters for each of the techniques. In addition, we experimented with the protein pocket representation to understand how the size and shape of a pocket impact final performance. The tested parameters and pocket setups are detailed in Appendix B.

### ArtiDock model

#### Feature extraction

The ligands were represented as atom-level molecular graphs. The node features included one-hot encoded atom group and the period from the periodic table. The graph edges represented covalent bonds between heavy atoms and the one-hot encoded covalent bond type.

Protein pocket node features were extracted for each heavy atom and included scalar (one-hot encoded atom group and period indexes, residue and atom names) and vector (distance from a pocket centroid to an atom) components. The graph edges were formed between a node and its 30 closest neighbors. The edge scalar features represented positional embedding calculated by the distance radial basis function [42]. The vectors between the nodes connected by edges were considered edge vector features.

#### Architecture

The ArtiDock is based on a proprietary model architecture inspired by the lightweight Trigonometry-Aware Neural Networks [4]. The model was built with an open-source machine learning framework, PyTorch v.2.5.1 [43], and a Graph Neural Network (GNN) library, PyTorch Geometric v.2.6.1 [44].

The ligand and pocket atom-level graphs were encoded using GNNs. The ligand graph was embedded by the graph isomorphism operator [45] (modified to incorporate edge features in the PyTorch Geometric library). The pocket graph was passed through Geometric Vector Perceptron GNN (GVP-GNN) [46], [47], which demonstrated superior performance in a range of tasks involving learning from protein structure. GVP-GNN uses both scalar and vector graph features as input. The resulting scalar and vector outputs are invariant and equivariant, respectively, for any rotation or reflection of a protein pocket in 3D Euclidean space.

After the graph encoders, the pocket and ligand node embeddings were combined in the pocket-ligand intermolecular embedding *z* ∈ ℝ^*p*×*c*×*d*^, where p - number of pocket nodes; c - number of ligand nodes; d - embedding size.

The intermolecular embedding was updated by a stack of blocks, each consisting of:

- update by intramolecular distance embedding of pocket *z*^*p*^ ∈ ℝ^*p*×*p*×*d*^ and ligand *z*^*c*^ ∈ ℝ^*c*×*c*×*d*^;
- multi-head self-attention;
- non-linear transition by MLP.

In the end, intermolecular embedding was converted into a pocket-ligand distance matrix *DM*^*pc*^ ∈ ℝ^*p*×*c*×1^ by a linear transformation followed by normalization.

#### Inference pipeline

The model outputs the pocket-ligand intermolecular distance matrix *DM*^*pc*^ for a given pocket and small molecule ligand. Thus, an additional distance matrix-to-pose algorithm is needed to convert the matrix into ligand atom coordinates (the actual binding pose). ArtiDock utilizes an algorithm that first infers a 3D point cloud from the distance matrix and then aligns a ligand conformer to the generated cloud. The algorithm details are available in Appendix C.

#### Physics-based optimization

In an effort to maximise the PoseBusters validity of ArtiDock inference pipeline output, we considered two post-processing strategies for the quick physics-based local minimization of the ligand binding poses:

- Local energy optimization by Vina v.1.2.6 [41] for up to 500 steps using Broyden-Fletcher-Goldfarb-Shanno method [48].
- Universal force field (UFF) [49] minimization for up to 500 iterations using RDKit v.2024.9.5. All pocket atoms were frozen, and only the ligand molecule was allowed to move.

### ArtiDock training

#### Dataset preparation

We used the train split of the PLINDER dataset as the model training dataset. The raw dataset contains 309,140 protein-ligand systems from 76,901 PDB IDs, comprising 34,103 unique ligands as identified by their CCD codes. The number of PLPs is 344,033.

After the initial preprocessing and quality filtering (Appendix A), the training dataset consisted of 227,066 PLPs from 206,555 systems and 67,888 PDB IDs. The complexes contained 29,440 unique ligand CCD codes. 34% of PLPs contained the ligands annotated as cofactors. 38% of PLPs comprised the ligands that follow Lipinski’s rule of five [50].

It is worth noting that, according to PLINDER metadata, 9% of the filtered PLPs involve ligands covalently bound to their targets. However, these were retained in the dataset as they met the quality filter criteria.

Since we retained all the entities in the complexes, each PLP may contain other ligands, ions, and water as part of the ligand binding pocket. In the filtered data, 12% of PLPs contained other non-cofactor ligands in the pockets; 8% contained cofactors; 16% included at least one ion. 48% of ligands had at least one water bridge interaction.

#### Data clustering

Many molecular structures are naturally overrepresented in the PDB as popular drug targets, model proteins, common drug-like ligands, cofactors, etc. For instance, among PLINDER PLPs, 3.6% of binding sites are represented by the HIV envelope glycoprotein GP120. Moreover, nearly 18% of PLPs contain ligands with a nucleoside core (e.g., ATP, UDP, NAD).

To remove the training data redundancy, we clustered the PLPs by ligand structure, protein pocket sequence similarity, and protein-ligand interaction similarity as described in Appendix D. As a result, 227,066 quality-filtered PLPs were grouped into 40,148 clusters, nearly half of which were singletons.

#### Training setup

The learning objective and other training protocol details are available in Appendix D.

Only PLPs that passed the quality filters were used for model training. We also excluded all entries that shared a PDB ID with any structure in the PoseBusters dataset to enable potential additional benchmarking of the trained model.

For each training epoch, we randomly sampled only one PLP from each cluster. However, we observed that certain ligand CCD codes remained overrepresented, as they appeared in multiple binding modes across a wide range of proteins (e.g., HSR, ADP, HEM). Therefore, we randomly downsampled these to participate in at most 0.1% of sampled PLPs.

Additionally, we introduced pocket-based data augmentation during the model training as detailed in Appendix D.

## Results and discussion

### Classical docking setup and parametrization

#### Docking parameters

The AutoDock-CPU experiments (Supplementary Table S1) revealed two parameters with considerable impact on the docking time/quality tradeoff: maximum number of energy evaluations and number of independent genetic algorithm runs. Both parameters are directly proportional to the computational cost. Reducing the maximum number of energy evaluations from the default 2.5 million to 350,000 achieved near-peak performance at approximately 7-fold docking speedup. Doubling the default number of runs (10) considerably increased the fraction of predictions with RMSD < 2Å from 0.42 to 0.46 (RMSD median decreased by 0.2Å). As a result, we used up to 350,000 energy evaluations and 20 genetic algorithm runs for the final benchmarking of AutoDock-CPU on PLINDER tests.

The AutoDock-GPU experiments (Supplementary Table S2) demonstrated minor changes in both computational cost and accuracy when varying docking parameters. We relate it to the well-adjusted ligand-based heuristics to define the maximum number of energy evaluations and the early stopping algorithm implemented in the docking tool. Thus, AutoDock-GPU was launched with default parameters for further benchmarking.

Similarly to the AutoDock-GPU, Vina uses heuristics to define the maximum number of evaluations. Overriding the parameter with a predetermined value didn’t seem to improve performance in terms of speed-to-quality ratio (Supplementary Table S3). The docking exhaustiveness, which scales computational cost linearly (in a single process mode), reached near-peak accuracy at the value of 16 (default is 8). Consequently, we used an exhaustiveness of 16 for PLINDER benchmarking.

Among the three Glide precision modes (Appendix B), SP demonstrated the best performance (Supplementary Table S4). It produced 49% of predictions with an RMSD < 2Å, significantly outperforming the faster HTVS mode (35%) and closely approaching the 51% achieved by the nearly 5 times slower XP mode. However, it is important to note that in terms of lDDT-PLI, a metric for evaluating protein-ligand interaction fidelity, the XP mode significantly outperformed SP, with a median score of 0.65 compared to 0.55. Therefore, XP may be a more suitable option when computational cost is not a limiting factor. When using SP mode, increasing the number of poses to use in post-docking minimization noticeably enhances prediction quality, adding only minor time overhead. Raising the number of poses from the default value of 5 to 80 reduced the median RMSD from 2.1Å to 1.8Å and improved the median lDDT-PLI score from 0.55 to 0.66, which is more accurate than XP mode. Further increase of the parameter value didn’t impact results. Considering these observations, Glide was launched in SP mode with the number of poses in post-docking minimization equal to 80 for PLINDER benchmarking.

#### Pocket representations

Surprisingly, the ligand-aligned bounding box (LABB) representation, which incorporates prior knowledge about the shape of the ligand binding mode, did not outperform the simple cubic box in any of the classical docking methods tested. In fact, the cubic box demonstrated even better performance in AutoDock-CPU and Glide experiments (Figure 2, Supplementary Tables S1-S4).

**Figure 2.**
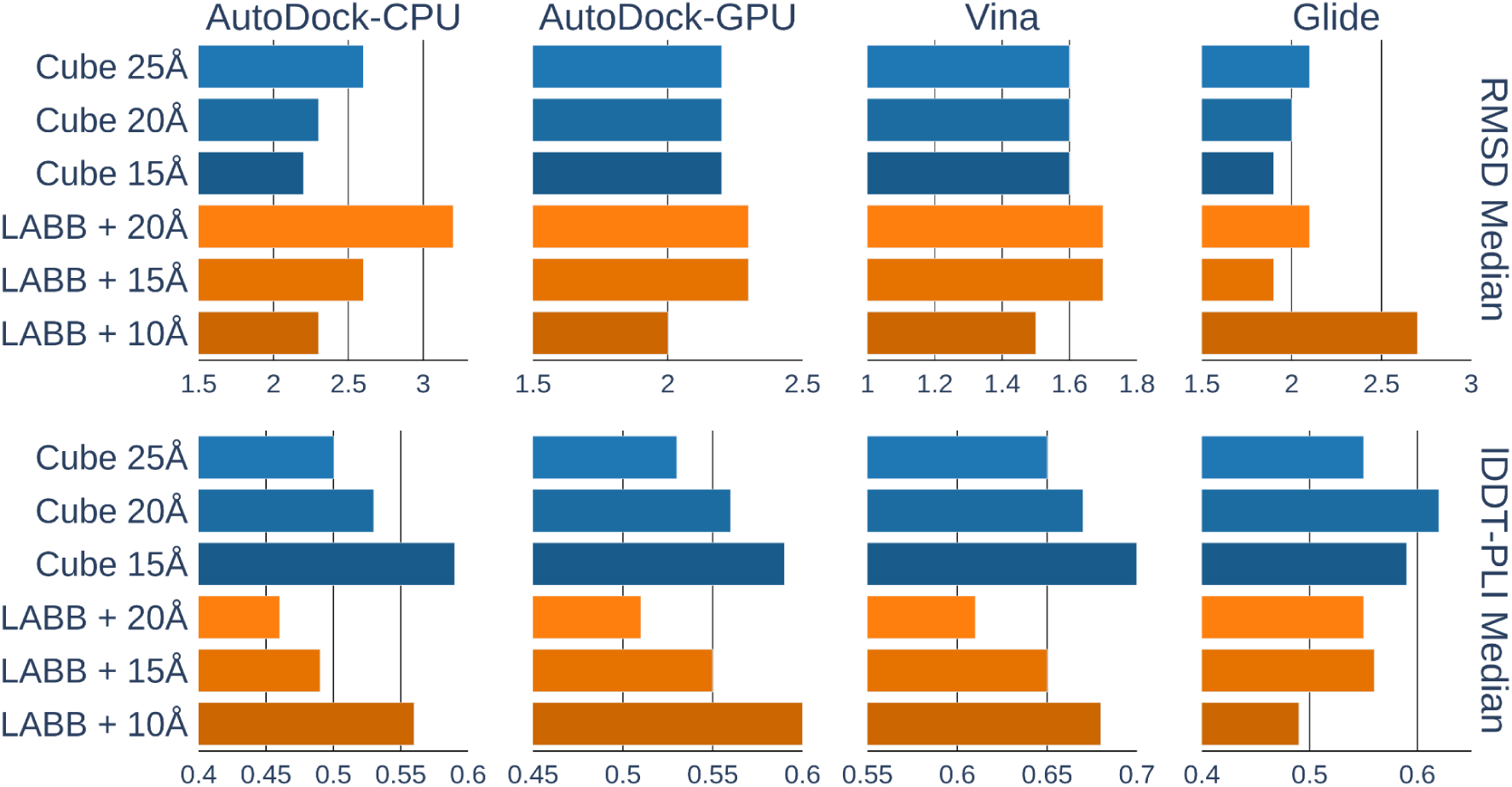
Classical docking top-ranked pose RMSD median (Å, smaller is better) and lDDT-PLI score median (larger is better) on PoseBusters dataset (N=306) when a cubic (Cube) box of varying sizes or a ligand-aligned bounding box (LABB) with varying dimension increments is used as pocket input.

Overall, reducing the docking box size either improved or did not affect accuracy across all tools, except for Glide, which achieved peak lDDT-PLI performance using a 20Å cubic box. This outcome is expected, as smaller pockets centered on the experimentally resolved ligand typically allow for a more exhaustive search in the correct region and reduce the likelihood of sampling highly-scored binding modes located far from the true binding site.

However, this highlights an important tradeoff between docking precision and the need for accurate pocket definition. In real-world applications, the exact position of the ligand is unknown, necessitating larger pockets to cover all plausible binding poses. To reflect this, we aimed to identify a shape-agnostic, sufficiently large pocket representation that still maintains high predictive accuracy for the final benchmarking. As a result, we selected the 20Å cubic box as the standardized input for all methods, including ArtiDock, in the PLINDER benchmark evaluation.

### PLINDER benchmark

The ArtiDock model, trained on the PLINDER training set to prevent data leakage, outperforms classical physics-based docking methods on the PLINDER test set (Table 1). The machine learning approach predicts ligand poses with a median RMSD of 2.0Å and a median lDDT-PLI score of 0.71, compared to 2.8Å and 0.64, respectively, for the best-performing traditional method, Glide.

**Table 1.** Performance of docking methods on the PLINDER test dataset (N=675). Green shading represents relative performance (darker is better).

| Method | RMSD Median (Å) | RMSD < 2Å | IDDT-PLI Median | IDDT-PLI > 0.5 | PB-Valid | RMSD < 2Å & PB-Valid | N Failed |
| --- | --- | --- | --- | --- | --- | --- | --- |
| AutoDock-CPU | 3.5 | 0.32 | 0.56 | 0.56 | 0.90 | 0.30 | 18 |
| AutoDock-GPU | 3.2 | 0.35 | 0.60 | 0.60 | 0.91 | 0.33 | 17 |
| Vina | 2.9 | 0.38 | 0.62 | 0.59 | **0.95** | 0.36 | 22 |
| Glide | 2.8 | 0.39 | 0.64 | 0.62 | 0.91 | 0.36 | 66 |
| ArtiDock | **2.0** | **0.48** | **0.71** | **0.79** | 0.72 | 0.38 | 0 |
| ArtiDock + Vina | 2.2 | 0.45 | 0.70 | 0.74 | 0.84 | **0.39** | 19 |
| ArtiDock + UFF | 2.3 | 0.41 | 0.69 | 0.76 | **0.95** | **0.39** | 14 |

Among the classical docking tools, Glide consistently delivers the best accuracy across most metrics, with Vina following closely behind. AutoDock-CPU, by contrast, shows the weakest performance with a median RMSD of 3.5Å.

As is evident from Figure 3, traditional docking methods exhibit significantly wider RMSD distributions than ArtiDock, with higher mean lDDT-PLI values (∼0.9 vs. ∼0.75). This indicates that they are occasionally able to predict very accurate poses, but at the same time produce significant amounts of low-quality ones. In contrast, ArtiDock produces many fewer low-quality predictions, resulting in better overall accuracy across the benchmark.

**Figure 3.**
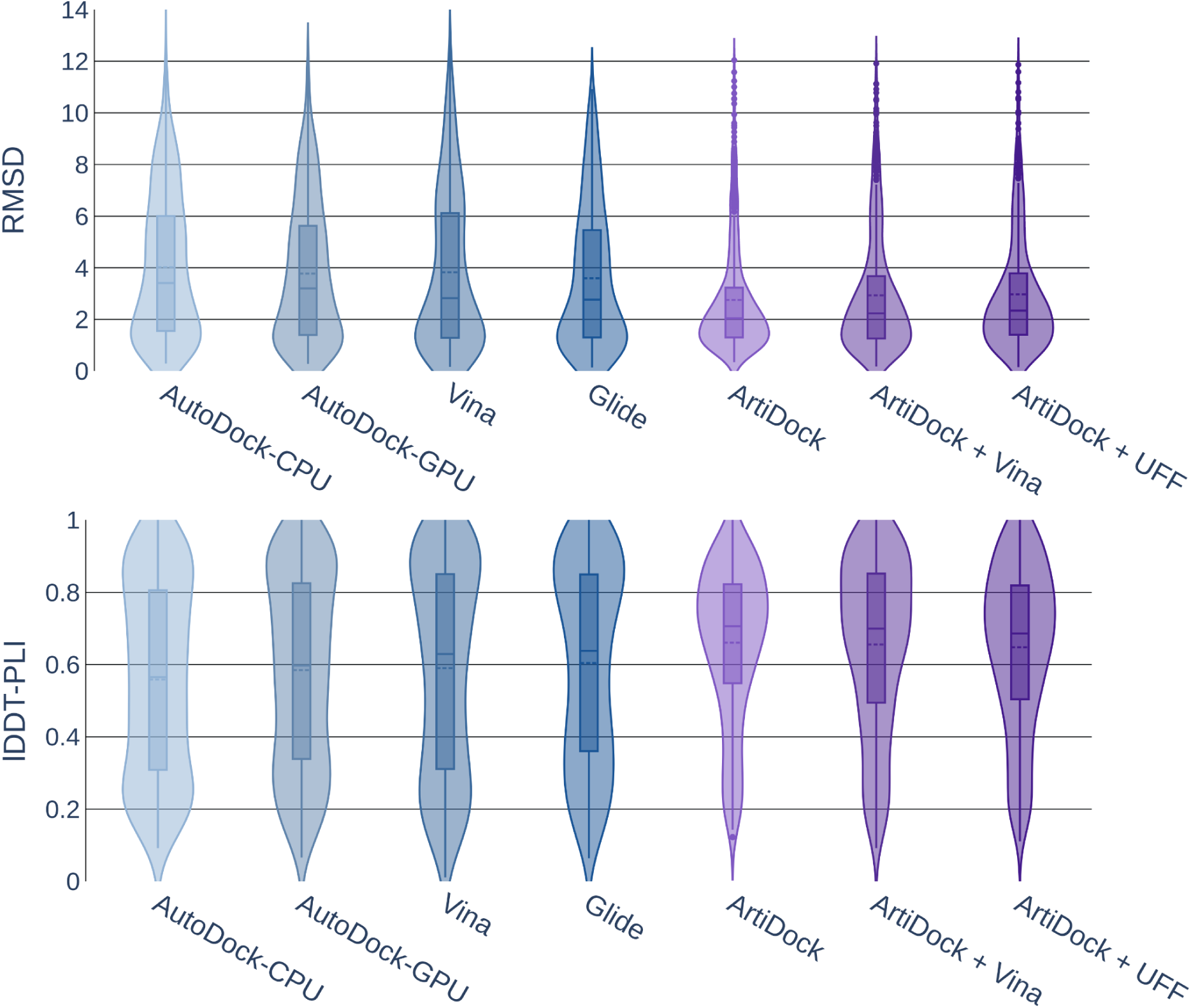
Violin plot of docking top-ranked pose RMSD (Å, smaller is better) and lDDT-PLI score (larger is better) distributions on the PLINDER test dataset (N=675). The bars represent the first and third quantiles, solid and dashed lines inside the bars indicate median and mean values, respectively. The shape represents the distribution of the values (kernel density plot).

The PoseBusters quality issues for all benchmarked tools predominantly originate from either too high internal energy of the predicted ligand pose or steric clashes with the pocket atoms (including heteroatoms), as shown in Supplementary Table S8. Considering physics-based approaches, the percentage of outputs with internal energy problems ranges from 5% in AutoDock-CPU/GPU to 1% in Glide. In contrast, Glide fails to avoid steric clashes in up to 7% of samples, while Vina produces clashes in only 1% of cases.

The fraction of ArtiDock predictions that pass the PB-Valid criterion is lower than that of the physics-based tools. However, applying a rapid post-prediction optimization using either Vina or RDKit UFF improves this metric at the cost of slightly lowering RMSD precision. Notably, UFF minimization increases the fraction of chemically and geometrically valid poses by 22%, matching Vina PB-Valid performance. Even after this correction, ArtiDock continues to outperform classical docking methods across all other reported metrics.

Estimated computational cost of ArtiDock is lower compared to the CPU-based classical tools and comparable to the AutoDock-GPU while providing much better accuracy (Appendix E).

We were unable to obtain any predictions for some PLPs, as shown in column “N Failed” of Table 1. Most failed predictions from AutoDock-CPU/GPU and Vina are due to the issues during protein or ligand preprocessing, for example, the inability to process molecules containing selenium or metal atoms. Of the 66 Glide failures, 11 are attributed to similar ligand preprocessing errors, while the remaining failures are primarily due to the absence of acceptable binding modes or the elimination of all poses during grid energy minimization. Failures during UFF minimization typically occur when the protein pocket cannot be handled correctly by RDKit.

It is important to note that we did not penalize any method for failed predictions. All reported results are based on the subset of successfully docked PLPs.

#### Docking with heteroatoms

The PLINDER structural data retains non-protein entities present in the original PDB structures, such as cofactors, ions, and ligand-bound water molecules (Appendix A). We included all heteroatoms within the input docking pocket and tracked docking performance across subsets of the data containing different types of these entities.

In general, the highest docking accuracy was observed in pockets containing organic cofactors (Figure 4), although this may be influenced by the relatively small sample size of just 23 PLPs.

**Figure 4.**
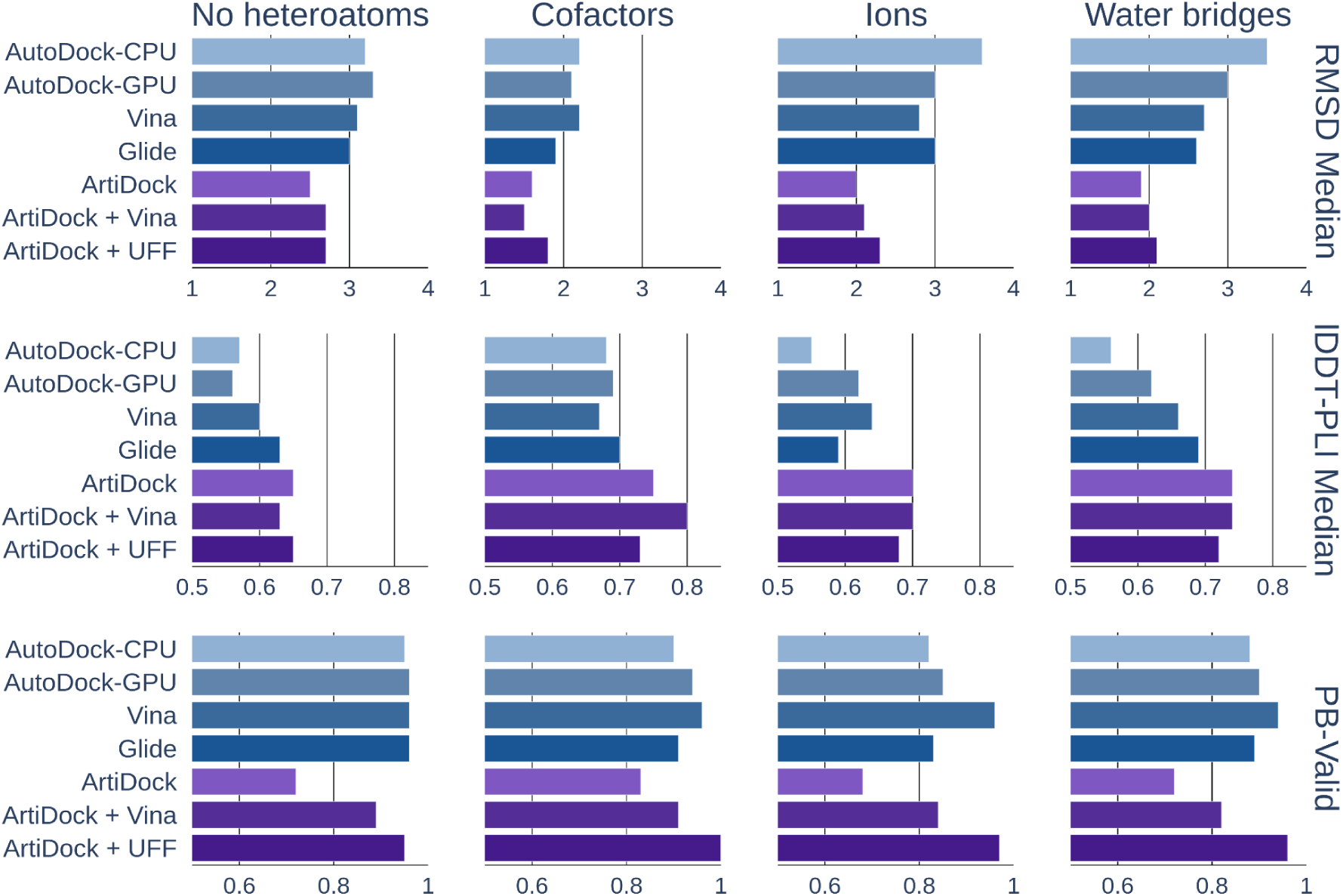
Docking top-ranked pose RMSD median (Å, smaller is better), lDDT-PLI score median (larger is better), and PB-Valid fraction (larger is better) on the PLINDER test data subsets with no heteroatoms (N=189), with cofactors (N=23), with ions (N=123), or ligand-bound water molecules (N=403) in a binding pocket.

ArtiDock consistently outperformed classical docking methods across all subsets, showing the greatest improvements in pockets containing ions and water. Specifically, ArtiDock achieved a median RMSD of 2.0Å in ion-containing pockets (vs. 2.8Å for the best classical method) and 1.9Å in water-containing pockets (vs. 2.6Å for the best classical method). According to PoseBusters’ quality checks, UFF-based optimization of ArtiDock predictions is on par with or outperforms the most successful classical approach, while Vina-based minimization yielded worse results.

Among physics-based tools, Glide provided the best precision in pockets containing cofactors, water, or protein only. Vina is the most accurate method in ion-containing pockets and shows the most reliable chemical validity across all subsets, with a fraction of PB-Valid results ranging from 0.94 to 0.96 (Figure 4).

The extended list of metrics is available in Supplementary Table S5.

### PLINDER MLSB benchmark

In contrast to the PLINDER test, which uses bound (holo) protein structures, the PLINDER MLSB subset is designed to evaluate docking tools under more realistic conditions, using apo or predicted protein inputs. This setting leads to a significant drop in performance, with most methods showing up as a twofold increase in median RMSD (Table 2).

**Table 2.** Performance of docking methods on the PLINDER MLSB test dataset (N=344). Green shading represents relative performance (darker is better).

| Method | RMSD<br>Median (Å) | RMSD < 2Å | IDDT-PLI<br>Median | IDDT-PLI ><br>0.5 | PB-Valid | RMSD < 2Å<br>& PB-Valid | N Failed |
| --- | --- | --- | --- | --- | --- | --- | --- |
| AutoDock-CPU | 5.9 | 0.10 | 0.30 | 0.23 | 0.91 | 0.10 | 6 |
| AutoDock-GPU | 6.2 | 0.09 | 0.28 | 0.21 | 0.91 | 0.09 | 5 |
| Vina | 6.5 | 0.08 | 0.27 | 0.17 | **0.94** | 0.07 | 5 |
| Glide | 5.5 | 0.13 | 0.33 | 0.26 | **0.94** | 0.13 | 54 |
| ArtiDock | **3.4** | **0.23** | **0.51** | **0.52** | 0.65 | **0.18** | 0 |
| ArtiDock + Vina | 4.2 | 0.19 | 0.46 | 0.43 | 0.80 | **0.18** | 5 |
| ArtiDock + UFF | 3.9 | 0.16 | 0.47 | 0.45 | 0.89 | 0.15 | 6 |

As in the holo benchmark, ArtiDock consistently outperforms all classical methods, achieving the lowest median RMSD (3.4Å) and highest median lDDT-PLI (0.51). The accuracy score distributions in the PLINDER MLSB test show that ArtiDock predicts a greater number of high-quality poses compared to other methods. In particular, ArtiDock’s lDDT-PLI score distribution peaks around 0.55, whereas Glide peaks at approximately 0.25, with even lower values for the other methods (Figure 5).

**Figure 5.**
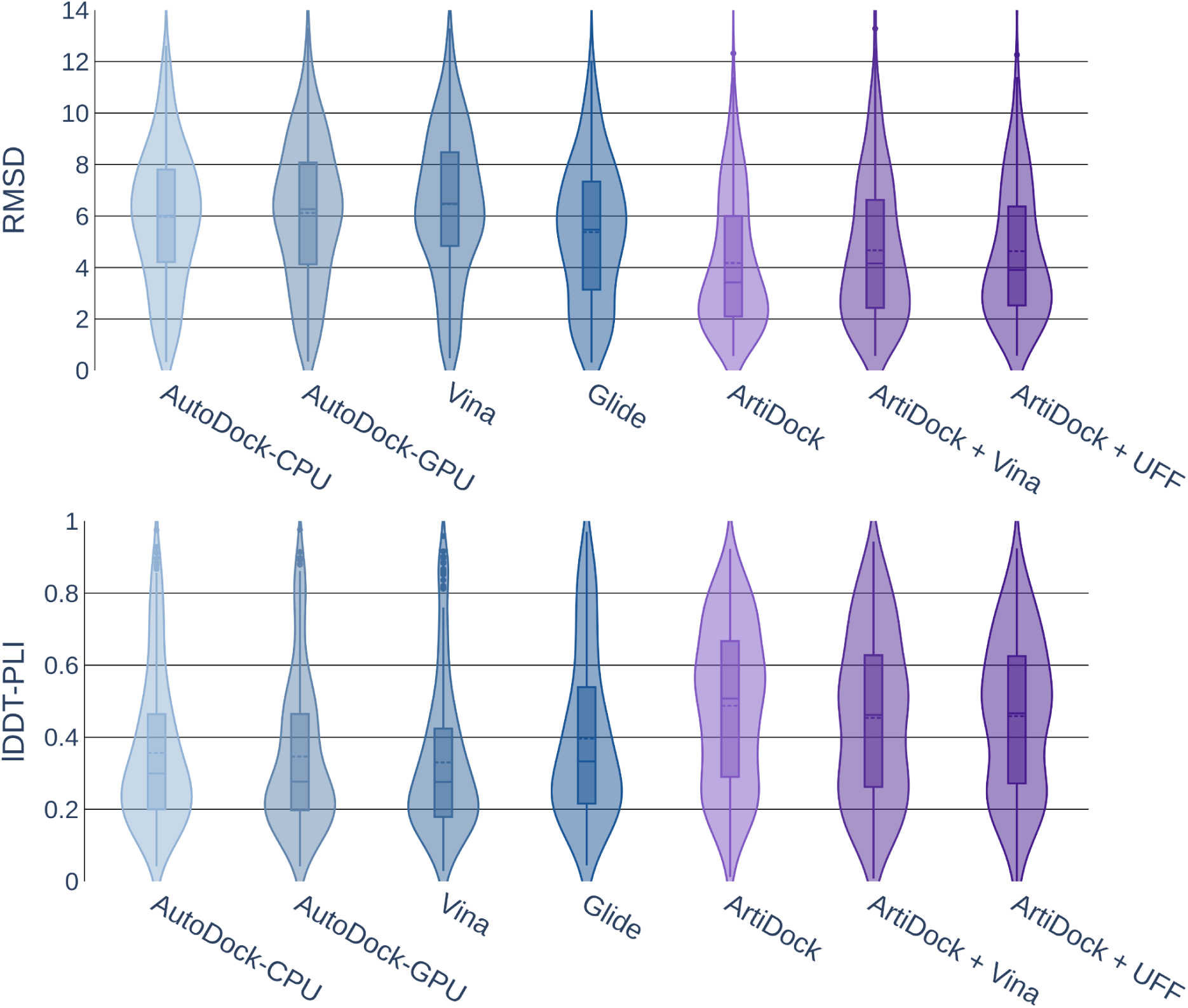
Violin plot of docking top-ranked pose RMSD (Å, smaller is better) and lDDT-PLI score (larger is better) distributions on the PLINDER MLSB test dataset (N=344). The bars represent the first and third quantiles, solid and dashed lines inside the bars indicate median and mean values, respectively (box plot). The shape represents the spread of values for a given method (kernel density plot).

Among the traditional tools, Glide maintains the best overall performance. Notably, Vina shows the steepest decline in accuracy, dropping from the second-best physics-based method in the PLINDER test to the worst performer in the PLINDER MLSB evaluation (Table 2). We partially attribute poor Vina performance on unbound structures to knowledge-based potentials used in the scoring function [24] that might cause some overfit to holo pocket representations.

The PB-Valid rate for ArtiDock in the MLSB test is up to 7 percentage points lower than in the holo test, regardless of post-prediction minimization. We attribute this decrease to cases where the ligand cannot physically fit into the unbound structure due to significant changes in pocket geometry. While classical docking methods tend to displace the ligand outside the binding box in response to steric clashes, ArtiDock predicts a distance matrix that may not fully account for such spatial incompatibilities but more accurately reflects the ground-truth binding mode.

Because ArtiDock inference pipeline prioritizes preserving predicted protein-ligand interatomic distances, the resulting poses in apo or predicted structures are more likely to exhibit steric issues. This trend is further supported by analysis involving the RMSD between superimposed Cα atoms in holo and apo/predicted pockets 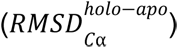. As shown in Supplementary Figure S1 and Supplementary Table S6, the ArtiDock non-minimized PB-Valid fraction decreases from 0.81 in the subset with 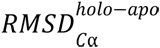 less than 0.1Å to 0.53 in the subset with 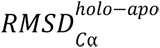 more than 0.5Å. In contrast, classical methods show no significant change in validity across these subsets.

When considering accuracy metrics, no correlation between 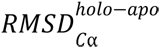 and either RMSD or lDDT-PLI was found (Supplementary Table S7). There was no noticeable decline in accuracy observed up to 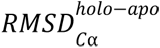 of 0.5Å. Nevertheless, larger deviations beyond this threshold result in a significant drop in RMSD and lDDT-PLI performance for all methods (Supplementary Figure S1, Supplementary Table S6). However, the part of PLPs with 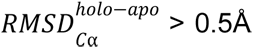 constitutes only 11.6%. To fully assess the effect of unbound structural deviations on docking performance, additional benchmarks with more structurally distinct apo/predicted proteins are needed.

## Limitations and perspectives

The current evaluation of ArtiDock was benchmarked against traditional docking tools on the PLINDER dataset. We ensured rigorous evaluation standards, which require retraining ML models on PLINDER training split. This limits our capacity to extensively benchmark alternative ML docking methods, especially heavyweight diffusion models and co-folding architectures. We invite other developers of ML-based docking tools to utilize and extend the PLINDER benchmark, thus contributing to a more comprehensive assessment of ML docking performance.

A broader range of classical docking tools, such as the commercially available CCDC GOLD [51] and DOCK [52], emerging open-source solutions like PandaDock [53], and various Vina-based alternatives [54], [55], [56], [57], should also be evaluated to cover the majority of physics-based docking techniques and to identify areas of improvement for their ML counterparts.

Chemical validity remains a concern in most ML-based docking approaches, including ArtiDock. Despite its high predictive accuracy, ArtiDock outputs often require additional physics-based post-processing steps to ensure chemical validity. Future efforts should be directed toward improving ArtiDock’s internal mechanisms to inherently produce chemically valid ligand poses, thus reducing or eliminating reliance on external physics-based adjustments.

Since there is no scoring function, which is assessed during the pose prediction, ArtiDock is not suitable for ranking alternative poses of the same ligand or comparing different ligands in terms of the binding strength, like conventional docking techniques do. The poses generated by ArtiDock are optimal for the given ligand and given conformation of the binding pocket, but their scoring should be performed externally by either algorithmic scoring functions or ML scoring models. To streamline and enhance predictive accuracy in high-throughput virtual screening scenarios, the development of a dedicated, ML-based scoring function tightly integrated within ArtiDock is necessary. Such an integrated scoring approach would likely provide enhanced applicability in real-world drug discovery applications.

## Conclusions

In this study, we presented ArtiDock, an ML-based approach for protein-ligand docking, which addresses the shortcomings of classical docking methodologies and is optimized to high-throughput virtual screening scenarios. The evaluation, conducted on the recently developed PLINDER benchmark dataset, demonstrates that ArtiDock predicts the ligand poses significantly more accurately in comparison to traditional physics-based docking techniques, reaching a median RMSD of 2.0Å and median lDDT-PLI of 0.71 compared to 2.8Å and 0.64, respectively, for the best-performing classical method, Glide.

ArtiDock’s lightweight model architecture and optimized inference pipeline allow it to reach the docking throughput, which is on par or better than the fastest conventional tools. This positions ArtiDock as a promising candidate for high-throughput virtual screening scenarios, overcoming one of the primary obstacles that have previously hindered widespread adoption of ML docking approaches.

ArtiDock is consistently more accurate than traditional techniques in various real-world docking scenarios, including pockets containing cofactors, ions, and water molecules. Notably, the pockets containing ions and water molecules showed especially pronounced accuracy gain, emphasizing ArtiDock’s robustness and adaptability to complex biologically relevant binding modes.

ArtiDock maintains a distinct advantage on the PLINDER MLSB dataset that includes especially challenging ligand-unbound protein structures, which deviate from the baseline ligand-bound conformations.

The integration of a quick, physics-based post-processing step substantially elevates the fraction of chemically valid poses to levels comparable to traditional docking approaches without compromising predictive accuracy or computational complexity significantly.

Overall, ArtiDock represents an advancement toward the much anticipated paradigm shift in docking towards ML-based tools, which effectively combines high throughput, robust predictive performance, and adaptability to realistic docking scenarios.

The future work should be primarily related to the inclusion of other classical and ML docking approaches to the current benchmark, the improvement of chemical and geometrical validity of ArtiDock output, and the implementation of ArtiDock’s native scoring function.

## Data and code availability

The source code for the classical docking setup, input data processing, result generation, and evaluation is available in the GitHub repository: https://github.com/receptor-ai/dock-eval.git.

The preprocessed PLINDER test subsets, predicted ligand poses from all compared tools, and tables with per-pose scores are stored in the Zenodo repository: https://zenodo.org/records/15493909.

The API of ArtiDock, trained on the PLINDER training split, is available by request for academic use. The ArtiDock trained on the full PDB data is commercially available at Receptor.AI.

## Author contributions

AN served as the conceptual lead and originator of the core idea. TV developed the ArtiDock model, inference pipeline, and docking parametrization module. IK and RS performed classical docking parameterization experiments, final evaluation, and analyzed results. TV and OK prepared PLINDER data and launched ArtiDock model training and evaluation. OK optimized ArtiDock inference pipeline. RK and IS researched and assessed existing methodologies of classical and ML-based molecular docking. SY and VH coordinated the work, critically assessed the results, and guided the analysis. SS and AN designed the study and controlled its progress. The manuscript was written by TV and SY.

## Conflicts of interest

All authors are employees of Receptor.AI INC. SS, AN, and SY have shares in Receptor.AI INC.

## Acknowledgements

SY has received funding through the grant MSMT-355/2025-16 from the Ministry of Education, Youth and Sport of the Czech Republic and through the MSCA4Ukraine project 101101923, which is funded by the European Union. Views and opinions expressed are, however, those of the author(s) only and do not necessarily reflect those of the European Union. Neither the European Union nor the MSCA4Ukraine Consortium as a whole nor any individual member institutions of the MSCA4Ukraine Consortium can be held responsible for them.

The authors acknowledge the use of the large language model GPT-4o for proofreading and stylistic editing of the manuscript.

## Notes

### Summary of Updates

The manuscript structure, images, and tables have fully changed. The ArtiDock model was trained and evaluated on another benchmark (PLINDER). The details of model inference were included in the Methods. Included a section on the parametrization of classical docking to reach optimal time/quality tradeoff. The model's performance was compared to parametrized classical docking.

## References

[1] X. Li, Y. Li, T. Cheng, Z. Liu, and R. Wang, “Evaluation of the performance of four molecular docking programs on a diverse set of protein-ligand complexes,” J. Comput. Chem., vol. 31, no. 11, pp. 2109–2125, Feb. 2010, doi: 10.1002/jcc.21498.

[2] J. B. Ghasemi, A. Abdolmaleki, and F. Shiri, “Molecular Docking Challenges and Limitations,” in Pharmaceutical Sciences: Breakthroughs in Research and Practice, IGI Global, 2017, pp. 770–794. doi: 10.4018/978-1-5225-1762-7.ch030.

[3] O. Méndez-Lucio, M. Ahmad, E. A. del Rio-Chanona, and J. K. Wegner, “A geometric deep learning approach to predict binding conformations of bioactive molecules,” Nat. Mach. Intell., vol. 3, no. 12, pp. 1033–1039, Dec. 2021, doi: 10.1038/s42256-021-00409-9.

[4] W. Lu, Q. Wu, J. Zhang, J. Rao, C. Li, and S. Zheng, “TANKBind: Trigonometry-Aware Neural NetworKs for Drug-Protein Binding Structure Prediction,” Jun. 06, 2022, Biophysics. doi: 10.1101/2022.06.06.495043.

[5] H. Stärk, O.-E. Ganea, L. Pattanaik, R. Barzilay, and T. Jaakkola, “EquiBind: Geometric Deep Learning for Drug Binding Structure Prediction,” Jun. 04, 2022, arXiv: arXiv:2202.05146. doi: 10.48550/arXiv.2202.05146.

[6] G. Zhou et al., “Uni-Mol: A Universal 3D Molecular Representation Learning Framework,” presented at the The Eleventh International Conference on Learning Representations, Sep 2022. Accessed: May 20, 2025. [Online]. Available: https://openreview.net/forum?id=6K2RM6wVqKu

[7] M. Buttenschoen, G. M. Morris, and C. M. Deane, “PoseBusters: AI-based docking methods fail to generate physically valid poses or generalise to novel sequences,” Chem. Sci., vol. 15, no. 9, pp. 3130–3139, 2024, doi: 10.1039/D3SC04185A.

[8] G. Corso, H. Stärk, B. Jing, R. Barzilay, and T. Jaakkola, “DiffDock: Diffusion Steps, Twists, and Turns for Molecular Docking,” Feb. 11, 2023, arXiv: arXiv:2210.01776. doi: 10.48550/arXiv.2210.01776.

[9] G. Corso, A. Deng, B. Fry, N. Polizzi, R. Barzilay, and T. Jaakkola, “Deep Confident Steps to New Pockets: Strategies for Docking Generalization,” Feb. 28, 2024, arXiv: arXiv:2402.18396.doi: 10.48550/arXiv.2402.18396.

[10] R. Krishna et al., “Generalized biomolecular modeling and design with RoseTTAFold All-Atom,” Science, vol. 384, no. 6693, p. eadl2528, Mar. 2024, doi: 10.1126/science.adl2528.

[11] Z. Qiao, et al., “NeuralPLexer3: Accurate Biomolecular Complex Structure Prediction with Flow Models,” Dec. 18, 2024, arXiv: arXiv:2412.10743. doi: 10.48550/arXiv.2412.10743.

[12] J. Abramson et al., “Accurate structure prediction of biomolecular interactions with AlphaFold 3,” Nature, vol. 630, no. 8016, pp. 493–500, Jun. 2024, doi: 10.1038/s41586-024-07487-w.

[13] C. Discovery et al., “Chai-1: Decoding the molecular interactions of life,” Oct. 15, 2024, bioRxiv. doi: 10.1101/2024.10.10.615955.

[14] J. Wohlwend, et al., “Boltz-1 Democratizing Biomolecular Interaction Modeling,” May 06, 2025, bioRxiv. doi: 10.1101/2024.11.19.624167.

[15] B. A. A. Team, et al., “Protenix - Advancing Structure Prediction Through a Comprehensive AlphaFold3 Reproduction,” Jan. 11, 2025, bioRxiv. doi: 10.1101/2025.01.08.631967.

[16] M. Leemann, A. Sagasta, J. Eberhardt, T. Schwede, X. Robin, and J. Durairaj, “Automated benchmarking of combined protein structure and ligand conformation prediction,” Proteins Struct. Funct. Bioinforma., vol. 91, no. 12, pp. 1912–1924, 2023, doi: 10.1002/prot.26605.

[17] P. Škrinjar, J. Eberhardt, J. Durairaj, and T. Schwede, “Have protein-ligand co-folding methods moved beyond memorisation?,” Feb. 10, 2025, bioRxiv. doi: 10.1101/2025.02.03.636309.

[18] J. Durairaj et al., PLINDER: The Protein-Ligand Interactions Dataset and Evaluation Resource. (Jul. 2024). Python. doi: 10.1101/2024.07.17.603955.

[19] H. M. Berman et al., “The Protein Data Bank,” Nucleic Acids Res., vol. 28, no. 1, pp. 235–242, Jan. 2000, doi: 10.1093/nar/28.1.235.

[20] “PLINDER Updates.” Accessed: May 17, 2025. [Online]. Available: https://www.plinder.sh/blog/updates

[21] G. M. Morris et al., “AutoDock4 and AutoDockTools4: Automated docking with selective receptor flexibility,” J. Comput. Chem., vol. 30, no. 16, pp. 2785–2791, 2009, doi: 10.1002/jcc.21256.

[22] D. S. Goodsell, M. F. Sanner, A. J. Olson, and S. Forli, “The AutoDock suite at 30,” Protein Sci., vol. 30, no. 1, pp. 31–43, 2021, doi: 10.1002/pro.3934.

[23] D. Santos-Martins, L. Solis-Vasquez, A. F. Tillack, M. F. Sanner, A. Koch, and S. Forli, “Accelerating AutoDock4 with GPUs and Gradient-Based Local Search,” J. Chem. Theory Comput., vol. 17, no. 2, pp. 1060–1073, Feb. 2021, doi: 10.1021/acs.jctc.0c01006.

[24] O. Trott and A. J. Olson, “AutoDock Vina: Improving the speed and accuracy of docking with a new scoring function, efficient optimization, and multithreading,” J. Comput. Chem., vol. 31, no. 2, pp. 455–461, Jan. 2010, doi: 10.1002/jcc.21334.

[25] J. Eberhardt, D. Santos-Martins, A. F. Tillack, and S. Forli, “AutoDock Vina 1.2.0: New Docking Methods, Expanded Force Field, and Python Bindings,” J. Chem. Inf. Model., vol. 61, no. 8, pp. 3891–3898, Aug. 2021, doi: 10.1021/acs.jcim.1c00203.

[26] R. A. Friesner et al., “Glide: A New Approach for Rapid, Accurate Docking and Scoring. 1. Method and Assessment of Docking Accuracy,” J. Med. Chem., vol. 47, no. 7, pp. 1739–1749, Mar. 2004, doi: 10.1021/jm0306430.

[27] T. A. Halgren et al., “Glide: A New Approach for Rapid, Accurate Docking and Scoring. 2. Enrichment Factors in Database Screening,” J. Med. Chem., vol. 47, no. 7, pp. 1750–1759, Mar. 2004, doi: 10.1021/jm030644s.

[28] R. A. Friesner et al., “Extra Precision Glide: Docking and Scoring Incorporating a Model of Hydrophobic Enclosure for Protein−Ligand Complexes,” J. Med. Chem., vol. 49, no. 21, pp. 6177–6196, Oct. 2006, doi: 10.1021/jm051256o.

[29] Y. Yang et al., “Efficient Exploration of Chemical Space with Docking and Deep Learning,” J. Chem. Theory Comput., vol. 17, no. 11, pp. 7106–7119, Nov. 2021, doi: 10.1021/acs.jctc.1c00810.

[30] A. N. Jain, A. E. Cleves, and P. Walters, “Deep-Learning Based Docking Methods: Fair Comparisons to Conventional Docking Workflows,” arXiv. doi: 2412.02889.

[31] “Machine Learning in Structural Biology.” Accessed: May 17, 2025. [Online]. Available: https://www.mlsb.io/

[32] S. Yesylevskyy, “MolAR: Memory-Safe Library for Analysis of MD Simulations Written in Rust,” J. Comput. Chem., vol. 46, no. 1, p. e27536, 2025, doi: 10.1002/jcc.27536.

[33] X. Robin, G. Studer, J. Durairaj, J. Eberhardt, T. Schwede, and W. P. Walters, “Assessment of protein–ligand complexes in CASP15,” Proteins Struct. Funct. Bioinforma., vol. 91, no. 12, pp. 1811–1821, Dec. 2023, doi: 10.1002/prot.26601.

[34] M. Biasini et al., “OpenStructure: an integrated software framework for computational structural biology,” Acta Crystallogr. D Biol. Crystallogr., vol. 69, no. 5, pp. 701–709, May 2013, doi: 10.1107/s0907444913007051.

[35] M. Bertoni, F. Kiefer, M. Biasini, L. Bordoli, and T. Schwede, “Modeling protein quaternary structure of homo- and hetero-oligomers beyond binary interactions by homology,” Sci. Rep., vol. 7, no. 1, Sep. 2017, doi: 10.1038/s41598-017-09654-8.

[36] forlilab/Meeko. (May 14, 2025). Python. Forli Lab. Accessed: May 15, 2025. [Online]. Available: https://github.com/forlilab/Meeko

[37] “MGLTools.” Accessed: May 19, 2025. [Online]. Available: https://ccsb.scripps.edu/mgltools/

[38] P. Karen, P. McArdle, and J. Takats, “Comprehensive definition of oxidation state (IUPAC Recommendations 2016),” Pure Appl. Chem., vol. 88, no. 8, pp. 831–839, Aug. 2016, doi: 10.1515/pac-2015-1204.

[39] “AutoDock.” Accessed: May 19, 2025. [Online]. Available: http://autodock.scripps.edu/

[40] ccsb-scripps/AutoDock-GPU. (May 15, 2025). C++. Center for Computational Structural Biology. Accessed: May 15, 2025. [Online]. Available: https://github.com/ccsb-scripps/AutoDock-GPU

[41] ccsb-scripps/AutoDock-Vina. (May 13, 2025). C++. Center for Computational Structural Biology. Accessed: May 15, 2025. [Online]. Available: https://github.com/ccsb-scripps/AutoDock-Vina

[42] J. Ingraham, V. Garg, R. Barzilay, and T. Jaakkola, “Generative Models for Graph-Based Protein Design,” in Advances in Neural Information Processing Systems, Curran Associates, Inc., 2019. Accessed: May 15, 2025. [Online]. Available: https://papers.nips.cc/paper_files/paper/2019/hash/f3a4ff4839c56a5f460c88cce3666a2b-Abstract.html

[43] A. Paszke, et al., “PyTorch: An Imperative Style, High-Performance Deep Learning Library,” 2019, arXiv. doi: 10.48550/ARXIV.1912.01703.

[44] M. Fey and J. E. Lenssen, “Fast Graph Representation Learning with PyTorch Geometric,” 2019, arXiv. doi: 10.48550/ARXIV.1903.02428.

[45] W. Hu, et al., “Strategies for Pre-training Graph Neural Networks,” 2019, arXiv. doi: 10.48550/ARXIV.1905.12265.

[46] B. Jing, S. Eismann, P. Suriana, R. J. L. Townshend, and R. Dror, “Learning from Protein Structure with Geometric Vector Perceptrons,” 2020, doi: 10.48550/ARXIV.2009.01411.

[47] B. Jing, S. Eismann, P. N. Soni, and R. O. Dror, “Equivariant Graph Neural Networks for 3D Macromolecular Structure,” 2021, arXiv. doi: 10.48550/ARXIV.2106.03843.

[48] J. Nocedal and S. J. Wright, Eds., “Sequential Quadratic Programming,” in Numerical Optimization, New York, NY: Springer, 1999, pp. 526–573. doi: 10.1007/0-387-22742-3_18.

[49] A. K. Rappe, C. J. Casewit, K. S. Colwell, W. A. Goddard, and W. M. Skiff, “UFF, a full periodic table force field for molecular mechanics and molecular dynamics simulations,” J. Am. Chem. Soc., vol. 114, no. 25, pp. 10024–10035, Dec. 1992, doi: 10.1021/ja00051a040.

[50] C. A. Lipinski, F. Lombardo, B. W. Dominy, and P. J. Feeney, “Experimental and computational approaches to estimate solubility and permeability in drug discovery and development settings,” Adv. Drug Deliv. Rev., vol. 46, no. 1–3, pp. 3–26, Mar. 2001, doi: 10.1016/S0169-409X(00)00129-0.

[51] M. L. Verdonk, J. C. Cole, M. J. Hartshorn, C. W. Murray, and R. D. Taylor, “Improved protein-ligand docking using GOLD,” Proteins, vol. 52, no. 4, pp. 609–623, Sep. 2003, doi: 10.1002/prot.10465.

[52] W. J. Allen et al., “DOCK 6: Impact of new features and current docking performance,” J. Comput. Chem., vol. 36, no. 15, pp. 1132–1156, 2015, doi: 10.1002/jcc.23905.

[53] P. K. Panda, pritampanda15/PandaDock. (May 20, 2025). Python. Accessed: May 21, 2025. [Online]. Available: https://github.com/pritampanda15/PandaDock

[54] A. Alhossary, S. D. Handoko, Y. Mu, and C.-K. Kwoh, “Fast, accurate, and reliable molecular docking with QuickVina 2,” Bioinformatics, vol. 31, no. 13, pp. 2214–2216, Jul. 2015, doi: 10.1093/bioinformatics/btv082.

[55] N. M. Hassan, A. A. Alhossary, Y. Mu, and C.-K. Kwoh, “Protein-Ligand Blind Docking Using QuickVina-W With Inter-Process Spatio-Temporal Integration,” Sci. Rep., vol. 7, no. 1, Art. no. 1, Nov. 2017, doi: 10.1038/s41598-017-15571-7.

[56] D. R. Koes, M. P. Baumgartner, and C. J. Camacho, “Lessons Learned in Empirical Scoring with smina from the CSAR 2011 Benchmarking Exercise,” J. Chem. Inf. Model., vol. 53, no. 8, pp. 1893–1904, Aug. 2013, doi: 10.1021/ci300604z.

[57] A. T. McNutt et al., “GNINA 1.0: molecular docking with deep learning,” J. Cheminformatics, vol. 13, no. 1, p. 43, Jun. 2021, doi: 10.1186/s13321-021-00522-2.

