## Appendices and Supplementary Figures for "ArtiDock: accurate Machine Learning approach to protein-ligand docking optimized for high-throughput virtual screening"

#### Supplementary Information

*Taras Voitsitskyi [1,2*], Ihor Koleiev [1,2], Roman Stratiichuk [1,3], Oleksandr Kot [1], Roman Kyrylenko [1], Illia Savchenko [1], Vladyslav Husak [1,4], Semen Yesylevskyy [1,2,5,6], Sergii Starosyla [1], Alan Nafiiev [1].*

[1] Receptor.AI Inc., 20-22 Wenlock Road, London N1 7GU, United Kingdom.

[2] Department of Physics of Biological Systems, Institute of Physics of The National Academy of Sciences of Ukraine, 46 Nauky Ave., 03038, Kyiv, Ukraine.

[3] Department of Biophysics and Medical Informatics, Educational and Scientific Centre "Institute of Biology and Medicine", Taras Shevchenko Kyiv National University, 64 Volodymyrska Str., 01601, Kyiv, Ukraine.

[4] Department of Cellular, Computational and Integrative Biology, The University of Trento, Via Sommarive 9, 38123 Povo (Trento), Italy.

[5] Institute of Organic Chemistry and Biochemistry, Czech Academy of Sciences, CZ-166 10 Prague 6, Czech Republic.

[6] Department of Physical Chemistry, Faculty of Science, Palacký University Olomouc, 17. listopadu 12, 771 46 Olomouc, Czech Republic.

*

#### Supplementary Figures

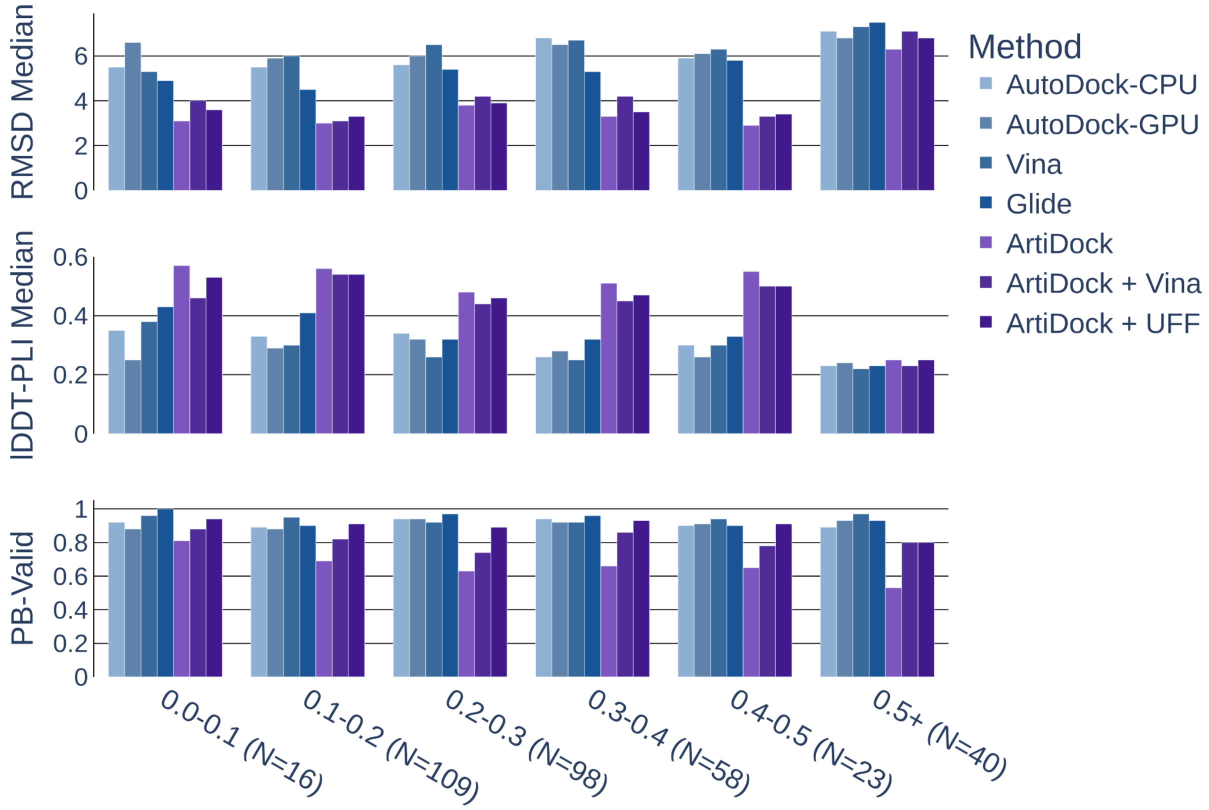


**Figure S1**. Docking top-ranked pose RMSD median (Å, smaller is better), lDDT-PLI score median (larger is better), and PB-Valid fraction (larger is better) on the PLINDER MLSB test data subsets at different Cα RMSD thresholds (Å) between superimposed holo and apo/predicted pockets.

### Appendix A. PLINDER data preparation

Each PLINDER system (complex) includes up to 5 proper ligand molecules, including cofactors. All the ions and experimental artifacts, as well as water molecules forming water bridges with any non-protein entity, are retained in the complex.

At the beginning of the dataset preprocessing, we removed all the hydrogen atoms from all the PLINDER system entities. Therefore, all the values related to atom counts or interatomic distances are reported for heavy (non-hydrogen) atoms only. We filtered out low-quality protein-ligand pairs (PLPs) by the following criteria:

- Base filters:
    - The number of protein residues is less than 5.
    - The number of ligand atoms is less than 5 or more than 100.
    - The number of rotatable bonds in the ligand is more than 50.
    - More than half of the ligand atoms are unresolved in the structure.

- Distance filters:
    - The minimum distance between ligand atoms and other entities in the complex is less than 1 Å. The distance is too low even for covalent bonds. Potential artifacts of the low-resolution structures or data representation errors.
    - The ratio of ligand steric clashes with other entities in the complex to the number of ligand atoms exceeds 0.1. The steric clash was defined as the distance between the atoms smaller than 70% of the sum of their respective van der Waals radii.
    - The average or standard deviation of the minimum distances from each ligand atom to other entities in the complex is more than 4.65 or 1.7, respectively. This filter helps to avoid complexes with a significant fraction of the ligand being too far away from the pocket. The threshold values were adjusted empirically by visual inspection of the complexes.
- Interaction filters
    - The number of non-covalent protein-ligand interactions is less than 3. The interactions are pre-computed in PLINDER metadata using the protein-ligand interaction profiler (PLIP) [1].
    - The ratio of non-covalent protein-ligand interactions to the number of ligand atoms is less than 0.1.

The extent of data loss due to quality filters is summarized in Table A1.

**Table A1**. Fraction of PLINDER PLPs passing quality filters.

| **Split** | **Base filters** | **Distance filters** | **Interaction filters** |
|---|---|---|---|
| **Train** | 0.99 | 0.82 | 0.75 |
| **Validation** | 0.99 | 0.89 | 0.95 |
| **Test** | 1.00 | 0.94 | 0.98 |

### Appendix B. Classical docking setup and parameterization details

#### Preprocessing issues

In the initial docking parameterization experiments on the PoseBusters dataset, we used MGLTools to prepare both proteins and ligands for AutoDock-CPU/GPU and converted docking output to the SDF format with Open Babel v.3.1.1 [2]. Meeko was used to process protein and ligand structures for Vina as suggested in the official documentation.

However, we observed that a substantial fraction of PLINDER test ligand poses generated using MGLTools + Open Babel were not readable by RDKit, which is essential for downstream evaluation. This issue was resolved by switching to Meeko for ligand handling. As for protein, the Meeko module design implies high-quality data with various structural issues being fixed manually, which is infeasible when working with large-scale datasets involving hundreds of proteins. Although excluding problematic residues or heteroatoms could improve compatibility, this would risk omitting components critical for ligand binding.

Therefore, MGLTools was prioritized over Meeko for the protein handling. Altogether, the combined use of Meeko and MGLTools allowed us to preserve up to 20% of predictions for further evaluation.

#### Docking parameters

The tested AutoDock-CPU genetic algorithm (GA) parameter space included:

- *ga_num_evals*: the maximum number of energy evaluations allowed during a GA run. This determines how long the search will continue. Once this limit is reached, the run stops regardless of whether convergence has occurred.
- *ga_run*: the number of independent GA runs to perform. Each run produces one predicted binding mode. More runs generally increase the chance of finding the global minimum.
- *ga_pop_size*: the number of solutions in the population during each generation.
- *ga_num_generations*: the maximum number of generations to be run per docking simulation.
- *ga_elitism*: the number of top individuals (best solutions) to survive to the next generation.

The GA parameters were changed in DPF before AutoDock-CPU docking.

In the AutoDock-GPU experiments, we tested the *nev*, *nrun*, *psize*, and *ngen* parameters that are equivalent to AutoDock-CPU *ga_num_evals*, *ga_run*, *ga_pop_size*, and *ga_num_generations,* respectively.

In the Vina experiments, we tested two main parameters responsible for the time/quality tradeoff: *exhaustiveness* (number of independent search attempts) and *max_evals* (number of scoring function evaluations).

In addition to the docking parameters, we examined whether grid resolution, defined by the *spacing* parameter, impacts the performance of AutoDock-CPU, AutoDock-GPU, and Vina.

For the Glide docking, we investigated such parameters as:

- *PRECISION*: one of *HTVS* (High Throughput Virtual Screening, high speed at the cost of accuracy), *SP* (Standard Precision, balance between speed and accuracy), or *XP* (Extra Precision, high accuracy at the cost of speed).
- *MAXKEEP*: number of binding modes to keep in the initial phase of docking.
- *MAXREF*: number of binding modes to keep for energy minimization.
- *MAX_ITERATIONS*: maximum number of conjugate gradient steps during the post-docking minimization.
- *POSTDOCK_NPOSE*: number of poses to use in post-docking minimization.

#### Pocket representations

The tested protein pocket size and shape representations included:

- Cube with side lengths equal to 15, 20, or 25 Å.
- Ligand-aligned bounding box (LABB) with side lengths equal to the maximal resolved ligand extent along each axis. We added 10, 15, or 20Å to each dimension for the experiments.

The pocket centroids were located at the center of the resolved ligand pose. The cube pocket represents a scenario when the exact binding site shape is not known or not important. The LABB implies some prior knowledge about ligand binding pose.

In the docking setup, pocket parameters were converted to the number of grid points during grid generation (GPF parameter *npts*) for AutoDock-CPU and AutoDock-GPU, directly passed as *box_size* parameter for Vina affinity maps generation, or directly passed as *OUTERBOX* parameter for Glide grid generation.

### Appendix C. ArtiDock inference pipeline details

For the point cloud generation, we combined and improved the ligand pose generation approaches from TankBind [3], [4], [5] and Uni-Mol [6]. Given the model predicted $DM^{pc}$, we zero-initialized the 3D point cloud $C' \in \mathbb{R}^{c\times 3}$ and applied back-propagation to optimize $C'$ with the Adam optimizer. The optimization stopped as soon as the loss reached a plateau. The loss function was calculated as the weighted sum of intramolecular and intermolecular distances and steric clash contributions.

The loss function included 4 components:

- Intermolecular distance loss:

$$\mathcal{L}_{inter} = \sum_{i}^{p}\sum_{j}^{c}\left(\left|DM_{ij}^{pc'} - DM_{ij}^{pc}\right|\right) \tag{1}$$

where $DM_{ij}^{pc'}$- distance matrix between pocket atoms and $C'$. This is the most fundamental component responsible for intermolecular atom positioning.

- Intramolecular distance loss:

$$\mathcal{L}_{intra} = \sum_{i}^{c}\sum_{j}^{c}\left(\left|DM_{ij}^{c'c'} - DM_{ij}^{cc}\right|\right) \tag{2}$$

for any ligand atom pair $ij \in LAS$, where LAS (local atomic structures) includes atom pairs connected by covalent bonds, or 2-hop away, or in the same ring structure [7], [8]; $DM_{ij}^{c'c'}$- intramolecular distance matrix calculated from $C'$; $DM_{ij}^{cc}$- ligand random conformer intramolecular distance matrix. It allows ligand flexibility during $C'$ optimization while preserving local intramolecular distance constraints.

- Intermolecular steric clash loss:

$$\mathcal{L}_{inter-clash} = \sum_{i}^{p}\sum_{j}^{c}\left(ReLU\left(VdW_{ij}^{pc} - DM_{ij}^{pc'}\right) + 1\right)^{2} \tag{3}$$

for any pocket atom i and ligand atom j if $VdW_{ij}^{pc} > DM_{ij}^{pc'}$, where ReLU - rectified linear unit function; $VdW^{pc} \in \mathbb{R}^{p \times c \times 1}$ - intermolecular sum of atom van der Waals radii multiplied by $T_{VdW} \in (0, 1)$; $T_{VdW}$ defines the fraction of sum of van der Waals radii below which a steric clash between atoms is formed. This loss was used to prevent pocket-ligand steric clashes.

- Intramolecular steric clash loss:

$$\mathcal{L}_{intra-clash} = \sum_{i}^{c}\sum_{j}^{c}\left(ReLU\left(VdW_{ij}^{cc} - DM_{ij}^{c'c'}\right) + 1\right)^{2} \tag{4}$$

for any ligand atom pair $ij$ if $VdW_{ij}^{cc} > DM_{ij}^{c'c'}$ and $ij \notin LAS$ and $i \neq j$, where $VdW^{cc} \in \mathbb{R}^{c \times c \times 1}$ - an intramolecular sum of atom van der Waals radii multiplied by $T_{VdW}$. Since potential distant intramolecular steric clashes were not accounted for in any of the previous losses, this component addresses the issue.

The final loss was calculated as the weighted sum of the abovementioned losses:

$$\mathcal{L} = w_{inter} * \mathcal{L}_{inter} + w_{intra} * \mathcal{L}_{intra} + w_{inter-clash} * \mathcal{L}_{inter-clash} + w_{intra-clash} * \mathcal{L}_{intra-clash} \quad (5)$$

where $w_{inter}$, $w_{intra}$, $w_{inter-clash}$, $w_{intra-clash}$ are corresponding loss weights.

Despite accounting for interatomic distance constraints during the distance matrix to point cloud transformation, the resulting point cloud $C'$ is still subject to artifacts and violations of geometric quality criteria. These issues are well known and were reported for the majority of ML-based docking methods [9].

We addressed the problem by applying the differential evolution (DE) technique as the final step of the distance matrix-to-pose algorithm. The DE is a stochastic approach that does not use gradient methods to find the minimum and can search large areas of the conformational space [10]. We utilized a modified version of DE for the ligand position optimization [11]. Given a point cloud $C'$ and a random input ligand conformer, we set the DE objective of RMSD minimization between $C'$ and the conformer coordinates by modifying the conformer rotatable bonds and aligning it to the $C'$. As a result, we obtain an optimized ligand conformer with atomic coordinates close to $C'$, which has significantly improved tetrahedral chirality, double bond stereochemistry, covalent bond lengths and angles, aromatic ring planarity, and double bond planarity.

To partially address the issue with steric clashes, we included additional inter- and intramolecular clash contributions to the DE objective function using conformer coordinates instead of $C'$ for defining the clashes.

Thus, the DE objective function was defined as follows:

$$\mathcal{L}^{DM} = RMSD + w^{DM}_{inter-clash} * \mathcal{L}^{DM}_{inter-clash} + w^{DM}_{intra-clash} * \mathcal{L}^{DM}_{intra-clash} \quad (6)$$

where $w^{DM}_{inter-clash}$ and $w^{DM}_{intra-clash}$ are corresponding loss weights.

The inference point cloud generation and DE objective function loss weights ($w_{inter}$, $w_{intra}$, $w_{inter-clash}$, $w_{intra-clash}$, $w^{DM}_{inter-clash}$, $w^{DM}_{intra-clash}$) together with $T_{VdW}$ and DE parameters were tuned using the PoseBusters quality checks on 300 unique ligand PLPs from the PLINDER validation data split. The tuning was performed for 200 trials with the help of parameters optimization software Optuna v.3.5.0 [12] using the tree-structured Parzen estimator algorithm. The best parameters were selected based on the highest fraction of the predicted ligand poses with RMSD < 2 Å and passing all the PoseBusters quality checks.

### Appendix D. ArtiDock training details

#### Data clustering

To address training data bias towards overrepresented PDB entities, we clustered PLPs according to:

- Ligand structure. We used Morgan fingerprints with a radius of 2 to assign each unique ligand to a cluster at the 0.5 distance threshold.
- Protein pocket sequence similarity between the systems. Clusters are derived from the community detection algorithm at a similarity threshold of 0.7. The clusters are precomputed in PLINDER metadata as the *pocket_fident_qcov__70__community* column.
- Protein-ligand interaction similarity between the systems. Clusters derived from the community detection algorithm at a similarity threshold of 0.7. The clusters are precomputed in PLINDER metadata as the *pli_qcov__70__community* column.

The clustering approaches and threshold values were selected empirically based on visual inspection of the entities grouped within each cluster. Preference was given to methods that produced a high number of representative clusters while minimizing the number of singleton clusters (clusters containing only one entry).

The final PLP clusters were defined as the union of the three previously described cluster types.

#### Training setup details

The training objective was based on the minimization of the difference between predicted and reference pocket-ligand intermolecular distance matrices using MSE loss.

The ground truth distance matrix was derived from the resolved ligand coordinates, while a randomly generated conformer of the full ligand was used as model input to obtain the predicted distance matrix. The distances were capped at 10Å to concentrate model capacity on predicting close-range interatomic contacts. During loss calculation, we accounted for the differences in ordering of ligand atoms between resolved and input structures, partially resolved ligands, and potential molecular symmetries.

We trained the model for 10 million iterations with the Adam optimizer. A batch size of 2 was used because of the GPU memory bottleneck. Subsequently, the Exponential Moving Average (EMA) technique was applied to smooth the noise in the training process and to improve the generalization of the model.

We used workstations with NVIDIA GeForce RTX 4090 GPU and AMD Ryzen 9 7950X3D 16-Core CPU for the ArtiDock model training

##### Augmented pocket representations

In each training iteration, a pocket was extracted randomly for a PLP, as all the protein residues/atoms:

- within a distance threshold from ligand atoms;
- inside a sphere/cube with a center in the ligand centroid;
- Inside a spheroid/cuboid reproducing ligand shape.

The distance thresholds and volume sizes were randomly sampled, with minimum and maximum values chosen to reflect the distribution of pocket atom counts obtained by including residues within 5 Å (minimum) to 8 Å (maximum) of the ligand. All the non-protein entities fitting into a pocket were retained.

The main idea behind the training on different pocket representations is to introduce to the model possible variations of the binding sites that might be used as input by the end user. In practice, a pocket is usually defined as a set of residues known/predicted to be in the binding site and/or residues/atoms around another ligand, whose binding pose is known. Subsequently, significant variability exists in potential pocket representations, which we address during training by augmenting pocket representations.

#### Appendix E. Estimating computational efficiency

A workstation with an AMD EPYC 9454P 48-Core Processor was employed to evaluate CPU-based docking methods (AutoDock-CPU, Vina, and Glide). The computations were parallelized using the Python multiprocessing module.

The computational cost of GPU-dependent methods was evaluated on the workstation with AMD Ryzen 9 7950X3D 16-Core Processor and NVIDIA GeForce RTX 3060 accelerated with CUDA 12.4.

Glide is the fastest CPU-based method, running about 3 times faster than AutoDock-CPU and 15 times faster than Vina. While Vina is inherently multithreaded, allowing it to reduce runtime for higher exhaustiveness settings when enough CPU cores are available, we ran it in a single-process mode and handled parallelization externally. This approach ensured consistent resource allocation across all CPU-based methods.

ArtiDock demonstrates comparable speed to another GPU-dependent method, AutoDock-GPU. In a virtual screening scenario, ArtiDock can reduce total runtime by overlapping GPU-based distance matrix prediction with CPU-based inference pipeline steps. ArtiDock's CPU-dependent post-processing is also parallelized efficiently and thus doesn't add much to the total runtime on modern multi-core machines.

For classical docking tools, the computational cost reflects optimized parameter settings aimed at achieving near-peak performance with minimal runtime, as described earlier.

Based on the average runtime per PLINDER test ligand and the hourly cost of the workstations used, we roughly estimate that docking one million small molecules would cost approximately $12 using GPU-based methods. This is followed by Glide at $15, AutoDock-CPU at $47, and Vina at $248. It is necessary to note that the actual costs may vary depending on multiple factors such as hardware configuration, pricing plan of the provider, pocket size, and ligands size and complexity.

### References


[1] M. F. Adasme *et al.*, "PLIP 2021: expanding the scope of the protein–ligand interaction profiler to DNA and RNA," *Nucleic Acids Res.*, vol. 49, no. W1, pp. W530–W534, Jul. 2021, doi: 10.1093/nar/gkab294.

[2] N. M. O'Boyle, M. Banck, C. A. James, C. Morley, T. Vandermeersch, and G. R. Hutchison, "Open Babel: An open chemical toolbox," *J. Cheminformatics*, vol. 3, no. 1, p. 33, Oct. 2011, doi: 10.1186/1758-2946-3-33.

[3] W. Lu, Q. Wu, J. Zhang, J. Rao, C. Li, and S. Zheng, "TANKBind: Trigonometry-Aware Neural NetworKs for Drug-Protein Binding Structure Prediction," Jun. 06, 2022, *Biophysics*. doi: 10.1101/2022.06.06.495043.

[4] M. R. Masters, A. H. Mahmoud, Y. Wei, and M. A. Lill, "Deep Learning Model for Efficient Protein–Ligand Docking with Implicit Side-Chain Flexibility," *J. Chem. Inf. Model.*, vol. 63, no. 6, pp. 1695–1707, Mar. 2023, doi: 10.1021/acs.jcim.2c01436.

[5] Z. Zsoldos, D. Reid, A. Simon, S. B. Sadjad, and A. P. Johnson, "eHiTS: A new fast, exhaustive flexible ligand docking system," *J. Mol. Graph. Model.*, vol. 26, no. 1, pp. 198–212, Jul. 2007, doi: 10.1016/j.jmgm.2006.06.002.

[6] G. Zhou *et al.*, "Uni-Mol: A Universal 3D Molecular Representation Learning Framework," May 26, 2022, *Chemistry*. doi: 10.26434/chemrxiv-2022-jjm0j.

[7] O. Trott and A. J. Olson, "AutoDock Vina: Improving the speed and accuracy of docking with a new scoring function, efficient optimization, and multithreading," *J. Comput. Chem.*, vol. 31, no. 2, pp. 455–461, Jan. 2010, doi: 10.1002/jcc.21334.

[8] H. Stärk, O.-E. Ganea, L. Pattanaik, R. Barzilay, and T. Jaakkola, "EquiBind: Geometric Deep Learning for Drug Binding Structure Prediction," Jun. 04, 2022, *arXiv*: arXiv:2202.05146. doi: 10.48550/arXiv.2202.05146.

[9] M. Buttenschoen, G. M. Morris, and C. M. Deane, "PoseBusters: AI-based docking methods fail to generate physically valid poses or generalise to novel sequences," *Chem. Sci.*, vol. 15, no. 9, pp. 3130–3139, 2024, doi: 10.1039/D3SC04185A.

[10] R. Storn and K. Price, "Differential Evolution – A Simple and Efficient Heuristic for global Optimization over Continuous Spaces," *J. Glob. Optim.*, vol. 11, no. 4, pp. 341–359, Dec. 1997, doi: 10.1023/A:1008202821328.

[11] O. M. Lucio, M. Ahmad, E. A. D. Rio-Chanona, and J. K. Wegner, "A Geometric Deep Learning Approach to Predict Binding Conformations of Bioactive Molecules (Dataset)," 2021, *figshare*. doi: 10.6084/M9.FIGSHARE.C.5407329.

[12] T. Akiba, S. Sano, T. Yanase, T. Ohta, and M. Koyama, “Optuna: A Next-generation Hyperparameter Optimization Framework,” 2019, *arXiv*. doi: 10.48550/ARXIV.1907.10902.
